# ATN status in amnestic and non-amnestic Alzheimer’s disease and frontotemporal lobar degeneration

**DOI:** 10.1101/2019.12.18.881441

**Authors:** Katheryn A.Q. Cousins, David J. Irwin, David A. Wolk, Edward B. Lee, Leslie M.J. Shaw, John Q. Trojanowski, Fulvio Da Re, Garrett S. Gibbons, Murray Grossman, Jeffrey S. Phillips

## Abstract

Under the ATN framework, cerebrospinal fluid analytes provide evidence of the presence or absence of Alzheimer’s disease pathological hallmarks: amyloid plaques (A), phosphorylated tau (T), and accompanying neurodegeneration (N). Still, differences in cerebrospinal fluid levels across amnestic and non-amnestic variants or due to co-occurring pathologies might lead to misdiagnoses. We assess the diagnostic accuracy of cerebrospinal fluid markers for amyloid, tau, and neurodegeneration in an autopsy cohort of 118 Alzheimer’s disease patients (98 amnestic; 20 non-amnestic) and 64 frontotemporal lobar degeneration patients (five amnestic; 59 non-amnestic). We calculated between-group differences in cerebrospinal fluid concentrations of amyloid-β_1–42_ peptide, tau protein phosphorylated at threonine 181, total tau, and the ratio of phosphorylated tau to amyloid-β_1–42_. Results show that non-amnestic Alzheimer’s disease patients were less likely to be correctly classified under the ATN framework using independent, published biomarker cutoffs for positivity. Amyloid-β_1–42_ did not differ between amnestic and non-amnestic Alzheimer’s disease, and receiver operating characteristic curve analyses indicated that amyloid-β_1–42_ was equally effective in discriminating both groups from frontotemporal lobar degeneration. However, cerebrospinal fluid concentrations of phosphorylated tau, total tau, and the ratio of phosphorylated tau to amyloid-β_1–42_ were significantly lower in non-amnestic compared to amnestic Alzheimer’s disease patients. Receiver operating characteristic curve analyses for these markers showed reduced area under the curve when discriminating non-amnestic Alzheimer’s disease from frontotemporal lobar degeneration, compared to discrimination of amnestic Alzheimer’s disease from frontotemporal lobar degeneration. In addition, the ATN framework was relatively insensitive to frontotemporal lobar degeneration, and these patients were likely to be classified as having normal biomarkers or biomarkers suggestive of primary Alzheimer’s disease pathology. We conclude that amyloid-β_1–42_ maintains high sensitivity to A status, although with lower specificity, and this single biomarker provides better sensitivity to non-amnestic Alzheimer’s disease than either the ATN framework or the phosphorylated-tau/amyloid-β_1–42_ ratio. In contrast, T and N status biomarkers differed between amnestic and non-amnestic Alzheimer’s disease; standard cutoffs for phosphorylated tau and total tau may thus result in misclassifications for non-amnestic Alzheimer’s patients. Consideration of clinical syndrome may help improve the accuracy of ATN designations for identifying true non-amnestic Alzheimer’s disease.

**Abbreviated Summary:** Cousins et al. assess the 2018 ATN framework and find that non-amnestic patients with Alzheimer’s disease (AD) have lower cerebrospinal fluid (CSF) phosphorylated tau and total tau than amnestic AD, while CSF amyloid-β accurately stratifies both non-amnestic and amnestic AD from frontotemporal lobar degeneration.

## Introduction

The 2018 ATN research framework is a systematic method to determine Alzheimer’s disease (AD) continuum designation, and can be applied using cerebrospinal fluid (CSF) analytes as markers of AD pathology (Jack *et al.*, 2018). Low CSF levels of amyloid-β 1–42 peptide (Aβ_1–_42) are associated with amyloid deposition in the brain, while high CSF levels of tau protein phosphorylated at threonine 181 (p-tau) are associated with intracellular hyperphosphorylated tau aggregation (Andreasen *et al.*, 2003; Tapiola *et al.*, 2009; Hampel *et al.*, 2018). Measures of CSF total tau (t-tau) have been interpreted as a marker of neurodegeneration that is not specific to AD; within the ATN framework, CSF t-tau can serve as a marker of disease severity (Jack *et al.*, 2016). Cut-points for CSF Aβ_1–42_, p-tau, and t-tau determine if patients should be judged as positive or negative for pathologic amyloid (A status), phosphorylated tau (T status), and neurodegeneration (N status), respectively (Jack *et al.*, 2018); these cut-points have been established in large autopsy cohorts (Shaw *et al.*, 2009). Under the ATN framework, A status determines whether an individual is positive or negative for Alzheimer’s continuum disease, while more fine-grained designations are determined by T and N status. Designations within Alzheimer’s continuum disease are based on the biological definition of AD that includes both amyloid plaques and tau tangles, and observations that changes in CSF Aβ_1–42_ typically precede CSF p-tau, followed by neurodegeneration and cognitive decline (Jack *et al.*, 2013). Thus, abnormal A and T markers (A+T+N- or A+T+N+) are interpreted as AD, while abnormal A alone (A+T-N-) is interpreted as early stage Alzheimer’s continuum or “Alzheimer’s pathologic change” and abnormal A and N (A+T-N+) is suspected co-pathology or “Alzheimer’s and suspected non-Alzheimer’s pathologic change”. Individuals who are A-negative are interpreted as not having Alzheimer’s continuum disease, and can have either normal biomarkers (A-T-N-) or biomarkers indicative of non-AD pathologic change (A-T+N-, A-T-N+, or A-T+N+). In this way, ATN provides a framework to interpret CSF biomarkers and to obtain a diagnosis in life beyond binary AD or non-AD.

Diagnostic sensitivity and specificity to typical, amnestic AD is high for CSF Aβ_1–42_, p-tau, and t-tau (Shaw *et al.*, 2009; Ewers *et al.*, 2015; Palmqvist *et al.*, 2015; Hampel *et al.*, 2018), and studies have shown that the ratio of p-tau to Aβ_1–42_ (p-tau/Aβ_1–42_) is especially accurate at stratifying patients with AD pathology from another form of neurodegenerative disease, frontotemporal lobar degeneration (FTLD) (Struyfs *et al.*, 2015; Oeckl *et al.*, 2016; Lleó *et al.*, 2018). Still, these studies demonstrate partial overlap in CSF levels between pathologic AD and FTLD. Thus, cut-points that determine ATN status still lead to a small but consequential subset of false-negative AD and false-positive FTLD cases. Such misdiagnoses can interfere with enacting appropriate clinical management strategies for patients and their families, and hinder the development of new therapeutic treatments and strategies by adding unexplained variance in a research population.

One possible source of diagnostic error is that broadly-applied cutoffs for AD pathology may better capture amnestic than non-amnestic variants of AD due to differences in CSF levels (Teng *et al.*, 2014; Paterson *et al.*, 2015; Wellington *et al.*, 2018; Pillai *et al.*, 2019). The majority of patients with AD pathology clinically present with amnestic AD, the most common form of dementia, characterized by profound loss of episodic memory (Dubois *et al.*, 2007). However, there are several non-amnestic variants of AD that instead present with visuospatial, language, or behavioral/executive impairments (Galton *et al.*, 2000; Murray *et al.*, 2011; Dickerson *et al.*, 2017). Patients with non-amnestic AD can phenotypically mimic and are often misdiagnosed as a clinical variant of frontotemporal dementia (FTD) (Koedam *et al.*, 2010), the second most common dementia. FTD is associated with pathologic FTLD (Hodges *et al.*, 2004; Perry and Miller, 2013) and, like non-amnestic AD, is characterized by language, behavioral, executive, and/or visuospatial dysfunction with relatively spared episodic memory. Both FTD and non-amnestic AD are characterized by a younger age of symptom onset than typical, amnestic AD (Koedam *et al.*, 2010; Crutch *et al.*, 2012; Lam *et al.*, 2013; Onyike and Diehl-Schmid, 2013; Mendez, 2017). Despite common AD pathology, amnestic and non-amnestic variants have relatively distinct patterns of anatomical disease distribution and protein accumulation (Galton *et al.*, 2000; Wolk, 2013, Phillips *et al.*, 2018*a, b*, 2019). Differences in disease dynamics between clinical syndromes may explain part of the observed variance in CSF levels amongst patients with AD pathology (Teng *et al.*, 2014; Paterson *et al.*, 2015).

Another potential source of diagnostic error is that CSF-derived ATN status may be relatively insensitive to primary pathologies other than AD. While the ATN framework interprets positive T and/or N status in the absence of A as non-AD pathology, these designations may not represent true A-negative status (Pouclet-Courtemanche *et al.*, 2019) or successfully detect other neurodegenerative pathologies, like FTLD. Moreover, co-occurring pathologies are common in persons with dementia, as well as in our brain bank (Robinson *et al.*, 2018). While an A+T-N+ profile is associated with Alzheimer’s and concomitant suspected non-Alzheimer’s pathologic change, it is unknown if this profile captures the majority of co-pathologic cases with primary non-AD pathology. Thus, secondary AD pathology — resulting in positive biomarkers for AD — may obscure a primary pathology of FTLD and reduce diagnostic specificity (Toledo *et al.*, 2012).

Here we used a sample of amnestic and non-amnestic patients with postmortem pathological diagnoses of primary AD or FTLD to test whether levels of CSF Aβ_1–42_, p-tau, and t-tau differ across amnestic and non-amnestic phenotypes, and whether these differences lead to errors in patients’ classification under the ATN framework. Similarly, we assessed pathologic and phenotypic differences for p-tau/Aβ_1–42_, a single biomarker that represents a commonly used alternative to the ATN framework for AD pathologic change and to stratify AD from FTLD (Struyfs *et al.*, 2015; Oeckl *et al.*, 2016; Lleó *et al.*, 2018). For each CSF analyte, we compared the frequency and clinical characteristics of true- and false-negative cases in non-amnestic and amnestic AD as well as true- and likely false-positives in FTLD. Receiver operating characteristic (ROC) curve assessments tested the diagnostic accuracy of CSF markers when stratifying patients with autopsy-confirmed primary AD from primary FTLD pathology. Between-group comparisons adjusted for age, sex, global cognitive impairment, APOE status, and presence of co-pathology. We compare A and T status during life to amyloid and tangle pathologic severity at autopsy, and we compare N status during life to postmortem atrophic severity. Our goals were to test the diagnostic accuracy of the ATN framework and the p-tau/Aβ_1–42_ ratio by comparing CSF markers across phenotypes and identifying likely sources of diagnostic error.

## Materials and methods

### Participants

Participants were 182 patients with a lumbar puncture and autopsy-confirmed primary AD or FTLD pathology identified through the Integrated Neurodegenerative Disease Biobank and Database (Xie *et al.*, 2011; Toledo *et al.*, 2014). All patients were autopsied at the University of Pennsylvania Center for Neurodegenerative Disease Research. Autopsy analyses classified the level of AD pathologic change from “not” to “high” using established “ABC” scoring methods (Montine *et al.*, 2012) for amyloid-β (modified Thal scoring on a four-point 0-3 scale), tau neurofibrillary tangles (modified Braak scoring on a four-point 0-3 scale), and neuritic plaques (four-point CERAD scale) when available (Mirra *et al.*, 1991). For FTLD-tau cases, Braak staging was determined using mAbs GT-7 and GT-38 immunohistochemistry that is specific to both 3 and 4 microtubule-binding repeat (3R; 4R) isoforms characteristic of AD neurofibrillary tangles (Gibbons *et al.*, 2018, 2019). Thirty cases lacked scoring for amyloid-β but had neurofibrillary tangle and CERAD neuritic plaque scores; cases with both CERAD and Braak scores of 2 or 3 were judged to have a non-zero likelihood of AD pathology, while cases with both CERAD and Braak scores of 0 or 1 were judged unlikely for AD pathologic change. For six cases, the level of AD pathologic change was unable to be estimated, and the presence of AD pathology was based on pathologists’ judgment. FTLD pathology was assessed using established histopathologic methods by accumulations of misfolded 3R or 4R tau (FTLD-Tau) associated with corticobasal degeneration, progressive supranuclear palsy, Pick’s disease, argyrophilic grain disease, and frontotemporal dementia with parkinsonism linked to chromosome 17; or transactive response DNA-binding protein of 43 kDa (TDP-43) associated with frontotemporal dementia and amyotrophic lateral sclerosis with cognitive impairment (FTLD-TDP) (Igaz *et al.*, 2008; Mackenzie *et al.*, 2010). Gross atrophic severity was visually evaluated by pathologists at autopsy on a four-point scale from none to severe.

Patients were followed clinically at the Penn Frontotemporal Degeneration Center or the Penn Memory Center and were clinically diagnosed by board-certified neurologists according to published diagnostic criteria (Gorno-Tempini *et al.*, 2011; McKhann *et al.*, 2011; Rascovsky *et al.*, 2011; Armstrong *et al.*, 2013; Höglinger *et al.*, 2017; Strong *et al.*, 2017). Participants’ consent was obtained according to the Declaration of Helsinki and approved by the University of Pennsylvania’s Institutional Review Board. Exclusion criteria included primary pathology other than AD or FTLD, patients without cognitive impairment, co-occurring neurologic conditions (e.g. stroke, hydrocephalus, chronic traumatic encephalopathy, vascular disease), and primary psychiatric disorders (e.g. depression, anxiety).

We next determined if AD and FTLD patients had mixed pathology, with primary pathological diagnosis based on pathologist’s judgment. AD patients were classified as having “mixed” pathology if they had α-synuclein positive Lewy bodies/neurites or TDP-43 inclusions (1-mild to 3-severe) in one or more neocortical regions (McKeith *et al.*, 2005; Montine *et al.*, 2012). FTLD patients were classified as having “mixed” pathology if they were positive for any level of AD pathology (ABC scores of low/intermediate/high) or neo-cortical α-synuclein (McKeith *et al.*, 2005; Montine *et al.*, 2012). Co-pathologic conditions included hippocampal sclerosis, limbic-predominant age-related TDP-43 encephalopathy (LATE), argyrophilic grain disease, Lewy body disease, primary age-related tauopathy (PART), and globular glial tauopathy. If no form of clinically meaningful co-pathology was detected, patients were classified as having “negligible” co-pathology. AD patients with amygdala-predominant Lewy bodies or cerebral amyloid angiopathy were considered to have “negligible” co-pathology. Of the 118 patients with confirmed primary AD pathology, 63 were classified as having mixed pathology. Three of these mixed cases had high levels of both FTLD and AD (high ABC rating), and primary pathology could not be determined; because of their high levels of AD pathologic change, these cases were classified as AD with mixed pathology in analyses. Of 64 patients with confirmed primary FTLD pathology, 34 were classified with mixed pathology. Of these FTLD with mixed pathology cases, three had ABC scores suggesting no AD, 22 had low AD pathologic change, and four had intermediate pathologic change. Of the remaining five FTLD cases missing ABC scores, co-pathology was determined by pathologists’ assessment; two were determined to have AD, one had argyrophilic grain disease, and two had PART.

Of 118 patients with confirmed AD pathology, 98 had an amnestic phenotype at presentation (clinical diagnoses: AD, n=95; amnestic mild cognitive impairment [aMCI], n=3), and 20 had a non-amnestic phenotype at presentation (clinical diagnoses: behavioral variant AD, [bvAD] n=1; behavioral variant FTD [bvFTD], n=2; unspecified FTD, n=5; unspecified primary progressive aphasia [PPA], n=2; logopenic variant PPA [lvPPA], n=3; corticobasal syndrome [CBS], n=7). Of 64 patients with confirmed FTLD pathology, 5 had an amnestic phenotype at presentation (clinical diagnoses: AD, n=5), and 59 had a non-amnestic phenotype at presentation (clinical diagnoses: non-amnestic MCI, n=1; bvFTD, n=16; unspecified FTD, n=8; unspecified PPA, n=3; lvPPA, n=3; nonfluent/agrammatic PPA, n=3; semantic variant PPA, n=5; CBS, n=9; progressive supranuclear palsy, n=7; and ALS with FTD, n=4).

### Demographic Comparisons

Across pathology and phenotype, we compared patients for number of copies of the APOE ε4 allele, sex, age at CSF, age at onset (earliest reported symptom), age at death, disease duration at CSF (years from onset to CSF), disease severity (Mini-Mental State Examination [MMSE]) (Folstein *et al.*, 1975), survival (years from earliest symptom onset to death), and years of education (Table 1). A chi-squared test indicated significant differences between AD and FTLD pathology in the frequency of individuals with zero, one, or two copies of the APOE ε4 allele (*Χ*^2^(2)= 35.70, *p=*1.77 × 10^−8^). Further testing indicated that the distribution of allele carriers did not differ between amnestic and non-amnestic AD groups (*Χ*^2^(2)=4.4, *p*=0.11), but that ε4 alleles were more frequent among the small group of amnestic FTLD patients than non-amnestic FTLD patients (*Χ*^2^(2)=14.78, *p*=0.0006). There was no significant difference in sex distribution across amnestic and non-amnestic AD and FTLD groups (*Χ*^2^(3)=1.17, *p*=0.76). Because quantitative demographic measures were not normally distributed, non-parametric Mann-Whitney-Wilcox tests compared AD and FTLD groups. AD and FTLD patients were not significantly different for years of education or disease duration (both *p*>0.1). AD patients were older at CSF compared to FTLD (W=5067.5, *p*=0.0001), and had marginally lower MMSE than FTLD (W = 2939.5, *p*=0.065). Kruskal-Wallis tests were used to compare across amnestic AD, non-amnestic AD, amnestic FTLD, and non-amnestic FTLD (Table 1). Patients were matched for MMSE, disease duration, education (all *p*>0.1), but were significantly different for age (*Χ*^2^(3)=19.86, *p*=0.0002). In pairwise comparisons using Mann-Whitney-Wilcox tests, amnestic AD patients were significantly older than non-amnestic FTLD (U=4108.5, *p=*1.02 x 10^−5^), and were marginally older than non-amnestic AD (U=1235, p=0.068). We observed similar differences for all other age-related variables (Table 1), including age at disease onset, age at death and survival, consistent with findings that non-amnestic variants tend to have an earlier disease onset than amnestic variants (Koedam *et al.*, 2010).

**Table 1:** Sample characteristics. Median (M), interquartile range (IQR), and sample size (n) by Pathology (Control, AD, FTLD) and Phenotype (amnestic, non-amnestic). For each variable, *X*^2^ and *p* values are reported for Chi-square or Kruskal-Wallis tests comparing the four patient groups (AD and FTLD by amnestic and non-amnestic).

|  |  | AD |  | FTL D |  | Chi-square/<br>Kruskal-Wallis |  |
| --- | --- | --- | --- | --- | --- | --- | --- |
| | | Amnestic | Non-amnestic | Amnestic | Non-amnestic | $\chi^2$ | $p$ |
| N<sub>total</sub> |  | 98 | 20 | 5 | 59 | — | — |
| Sex (% male) |  | 48% | 50% | 33% | 54% | 1.2 | 0.76 |
| APOE ε4<br>alleles | 0 | 32% | 50% | 50% | 87% | 47.1 | <0.001 |
|  | 1 | 49% | 44% | 33% | 13% |  |  |
|  | 2 | 19% | 6% | 17% | 0% |  |  |
| Age at | M | 73.5 | 63.5 | 71.0 | 65.0 | 19.9 | <0.001 |
| CSF | IQR | 14.5 | 13.5 | 4.0 | 10.0 |  |  |
| (years) | n | 98 | 20 | 5 | 59 |  |  |
| Duration at | M | 3.0 | 2.5 | 6.0 | 4.0 | 2.7 | 0.44 |
| CSF | IQR | 3 | 4 | 3 | 3 |  |  |
| (years) | n | 98 | 20 | 5 | 59 |  |  |
| Age at | M | 69 | 61 | 66 | 61 | 19.8 | <0.001 |
| Onset | IQR | 15.75 | 13.0 | 6.0 | 11.5 |  |  |
| (years) | n | 98 | 20 | 5 | 59 |  |  |
| Age at | M | 79.0 | 72.5 | 76.0 | 69.0 | 31.9 | <0.001 |
| Death | IQR | 15.0 | 11.5 | 8.0 | 10.0 |  |  |
| (years) | n | 98 | 20 | 5 | 59 |  |  |
| Survival | M | 9.0 | 8.5 | 9.0 | 7.0 | 9.5 | 0.02 |
| (years) | IQR | 5 | 6 | 1 | 5 |  |  |
|  | n | 98 | 20 | 5 | 59 |  |  |
| Education | M | 15.5 | 15.0 | 16.0 | 16.0 | 1.8 | 0.62 |
| (years) | IQR | 5.0 | 4.0 | 0.0 | 5.75 |  |  |
|  | n | 98 | 19 | 5 | 58 |  |  |
| MMSE | M | 23 | 19 | 24 | 25 | 5.6 | 0.13 |
| (max=30) | IQR | 9 | 9 | 8 | 7 |  |  |
|  | n | 97 | 17 | 5 | 57 |  |  |

### Cerebrospinal fluid (CSF) analysis

CSF samples were collected following a standard lumbar puncture typically following an overnight fast. Samples were measured for Aβ_1–42_, p-tau, t-tau, and p-tau/Aβ_1–42_ using the xMAP Luminex platform (INNO-BIA AlzBio3 for research-only reagents; Innogenetics). Analyses were performed by laboratory technicians who were blinded to clinical data. Published cut-points for all measures are based on best thresholds reported by Shaw and colleagues (2009); amyloid (A) positivity was based on an Aβ_1–42_ cutoff of 192 pg/mL; tau (T) positivity was based on a p-tau cutoff of 23 pg/mL; neurodegeneration (N) positivity was based on a t-tau cutoff of 93 pg/mL; and a cutoff of 0.10 was applied to the p-tau/Aβ_1–42_ ratio to determine positivity for AD. Outlier checks were performed on CSF values, and one case in the pathologic AD group was excluded for a CSF t-tau value more than five standard deviations above the sample mean.

### Statistical analysis

#### ATN Classification Based on CSF Biomarkers

For phenotypic and pathologic groups, we evaluated ATN designation of patients on the AD continuum based on positive or negative A, T, or N status and examined the profiles of cases that were likely misclassified. Patients with CSF Aβ_1–42_ ≤ 192 pg/mL were considered A-positive; those with CSF p-tau ≥ 23 pg/mL were considered T-positive; and those with CSF t-tau ≥ 93 pg/mL were considered N-positive. In addition, we tested classification using the p-tau/Aβ_1–42_ ratio; patients with p-tau/Aβ_1–42_ ≥ 0.10 were considered positive for AD.

#### Between-group comparisons of CSF

To elucidate the sources of misclassification within the ATN framework or the p-tau/ Aβ_1–42_ ratio, analyses of covariance (ANCOVAs) compared each CSF analyte across pathology (AD, FTLD) and phenotype (amnestic, non-amnestic), covarying for number of APOE ε4 alleles, co-pathology status (negligible, mixed), age at CSF, MMSE, and sex (α = 0.05). To ensure that the presence of mixed pathology was not driving differences, models were also run excluding patients with high levels of co-pathology. MMSE scores were unavailable for five participants (two non-amnestic AD, one amnestic AD, and two non-amnestic FTLD), and education data were unavailable for two participants (one non-amnestic AD and one non-amnestic FTLD). These missing data were imputed based on the mean of each patient’s respective pathology and phenotype group. In initial statistical analysis of CSF measures, model residuals were not normally distributed, violating an assumption of multivariate normality; thus, a log transformation was applied to each measure. Because of the unbalanced design, type II sum of squares were calculated for ANCOVAs. CSF levels were compared through post-hoc linear contrasts.

For each marker, true- and false-negative and true- and likely false-positive cases were identified based on established cut-points (Shaw *et al.*, 2009), and Fisher’s exact tests were used to test whether the likelihood of being classified as positive for each marker differed according to memory phenotype in AD. We did not compare CSF marker positivity between FTLD phenotypes due to the small number of amnestic FTLD cases (n=6).

#### Discrimination of pathologic AD from FTLD

Finally, receiver operating characteristic (ROC) curve and area under the curve (AUC) analyses assessed diagnostic accuracy of each CSF marker when discriminating non-amnestic patients with AD from FTLD pathology, compared to discriminating amnestic patients with AD from FTLD pathology. Optimal cutoffs were calculated for each CSF marker on the basis of Youden’s index, which maximized the sum of sensitivity and specificity in the current sample.

All statistical analyses were conducted in the R statistical environment, using the Companion to Applied Regression (car), multcomp, and partial ROC (pROC) packages (Robin *et al.*, 2011; Fox *et al.*, 2012; R Core Team, 2017).

### Data Availability

All qualified investigators are welcome to view data through an established algorithm at the CNDR.

## Results

### ATN Classification Based on CSF Biomarkers

Table 2 outlines patient classifications according to A, T, and N positivity based on published cutoffs for CSF biomarkers (Shaw *et al.*, 2009). According to the ATN framework, of the 98 amnestic patients with autopsy-confirmed AD pathology, 91 (93%) had CSF biomarkers positive for Alzheimer’s continuum disease. A similarly high proportion of non-amnestic patients with AD pathology, 17 of 20 (85%), were positive for Alzheimer’s continuum disease. However, the likelihood of more specific ATN designations was different across amnestic phenotype in patients with AD pathology. Fisher’s tests indicated that non-amnestic AD patients were significantly less likely to be positive for all three markers than amnestic AD patients (20% vs. 52%; OR = 0.23; CI = 0.05–0.79; *p*=0.013), and were significantly more likely to be negative for all three markers and classified as having normal biomarkers than amnestic AD patients (15% vs. 2%; OR = 8.23; CI = 0.88–105.4; *p*=0.034).

**Table 2:** Results of ATN classification using CSF biomarkers. Interpretation of biomarkers using ATN framework (above the black line) and p-tau/Aβ_1–42_ ratio (below the black line).

| Interpretation |  | Biomarker profile | Amnestic AD (n=98) | Non-amnestic AD (n=20) | Amnestic FTLN (n=5) | Non-amnestic FTLN (n=59) |
| --- | --- | --- | --- | --- | --- | --- |
| Alzheimer's continuum | Alzheimer's disease | A+T+N+ | 51 (52%) | 4 (20%) | 0 (0%) | 0 (0%) |
|  |  | A+T+N- | 16 (16%) | 5 (25%) | 0 (0%) | 0 (0%) |
|  | Alzheimer's pathologic change | A+T-N- | 13 (13%) | 6 (30%) | 2 (40%) | 7 (12%) |
|  | AD and concomitant suspected non-AD pathologic change | A+T-N+ | 11 (11%) | 2 (10%) | 1 (20%) | 2 (3%) |
| Non- Alzheimer's pathologic change |  | A-T+N- | 2 (2%) | 0 (0%) | 0 (0%) | 2 (3%) |
|  |  | A-T-N+ | 0 (0%) | 0 (0%) | 0 (0%) | 7 (12%) |
|  |  | A-T+N+ | 3 (3%) | 0 (0%) | 0 (0%) | 3 (5%) |
| Normal biomarkers |  | A-T-N- | 2 (2%) | 3 (15%) | 2 (40%) | 38 (64%) |
| Alzheimer's disease | | p-tau/A$\beta_{1-42} \geq 0.1$ | 88 (90%) | 13 (65%) | 1 (20%) | 9 (15%) |

Only ten of 118 AD cases (8%) were negative for Alzheimer’s continuum disease. Of these, five had profiles indicative of non-AD pathologic change (A-T+N- or A-T+N+); all five were amnestic; two had no co-occurring pathology detected at autopsy, two had co-occurring Lewy body pathology, and one had co-occurring TDP-43 pathology. ABC scores indicated high (n=2) or intermediate (n=3) AD pathologic change. Another five (three non-amnestic; 2 amnestic) had normal profiles (A-T-N-); ABC scores show high (n=3), intermediate (n=1), and low (n=1) levels of AD pathologic change; two patients had negligible co-pathology and three had co-occurring TDP-43 pathology.

We also examined accuracy of the A+T-N+ classification to indicate concomitant Alzheimer’s and non-Alzheimer’s pathologic change. Thirteen of 118 patients (11 amnestic; two non-amnestic) were classified as A+T-N+. ABC scores indicated high (n=12) or intermediate (n=1) AD pathologic change. Four of these 13 (31%) patients were found to have negligible levels of co-occurring pathology at autopsy and represent probable misclassifications.

Of the 64 FTLD patients, only 12 (19%; 12 non-amnestic; zero amnestic) had an ATN profile suggestive of non-AD pathologic change (A-T+N+, A-T-N+, or A-T+N-). The majority of FTLD patients (63%) were classified as having normal ATN biomarkers. Nine (14%) FTLD patients had an A+T-N-profile, suggesting AD pathologic change only. Of these nine, two were determine to not have AD (ABC scores, n=2; pathologist, n=1); these cases represent probable misclassifications. The other six were determined to have low AD pathologic change (ABC scores, n=5; pathologist, n=1); ATN biomarkers in these cases were insensitive to patients’ primary FTLD pathology. Three FTLD patients had an A+T-N+ profile, suggesting concomitant AD and non-AD pathologic change; two were found to have negligible co-pathology at autopsy, and the final patient had likely AD pathology.

Finally, we compared the ATN framework to the p-tau/Aβ_1–42_ ratio in amnestic and non-amnestic AD patients. Performance was equivalent in patients with autopsy-confirmed AD pathology. Of 98 amnestic AD patients, 91 (93%) were classified as positive for Alzheimer’s continuum using the ATN framework (as reported above), and 88 (90%) were classified as positive for AD using the p-tau/Aβ_1–42_ ratio. Of the 20 non-amnestic patients with AD pathology, 17 (85%) were classified as positive for Alzheimer’s continuum using the ATN framework (as reported above), while 13 (65%) were classified as positive for AD using the p-tau/Aβ_1–42_ ratio. Comparing the false-positives in FTLD patients with negligible co-pathology (Supplementary Table 1), two of 30 FTLD patients (7%) were positive for AD pathology using the p-tau/Aβ_1–42_ ratio, and four of 30 FTLD patients (13%) were positive for Alzheimer’s continuum disease according to the ATN framework. While ATN identified a numerically larger proportion of non-amnestic AD cases, and the p-tau/Aβ_1–42_ ratio had fewer false-positives, Fisher’s tests determined these proportions were not significantly different (all *p*>0.2).

Chi-squared tests indicated no significant differences between amnestic and non-amnestic patients with AD pathology in Thal, Braak, or CERAD staging (all *p*>0.3), nor in brain atrophy ratings between patients with FTLD and AD pathology (*p*>0.5). To elucidate the sources of ATN classification differences between non-amnestic and amnestic AD, we subsequently compared levels of each CSF marker parametrically across pathology and phenotype. If CSF Aβ_1–42_, p-tau, and the p-tau/Aβ_1–42_ ratio are accurate markers of likely amyloid, tangle, and AD pathology respectively, they are expected to be significantly different across pathology (AD, FTLD) but invariant to phenotype (amnestic, non-amnestic). If CSF t-tau is an accurate non-specific staging marker of N, all patient groups would be hypothesized to have a high proportion of N-positive cases; while we do not know severity of neurodegeneration in life, all patients were symptomatic at time of CSF sample (Jack *et al.*, 2013).

### Between-group comparisons of CSF Aβ_1–42_

We compared levels of CSF Aβ_1–42_ parametrically across pathology and phenotype (Figure 1). An ANCOVA (Type II) showed that Aβ_1–42_ levels differed by pathology (F(1,173)=56.2, *p=*3.2 × 10^−12^), with lower values in the pathologic AD group than the pathologic FTLD group. Additionally, Aβ_1–42_ levels were significantly associated with APOE status (F(1,173)=6.1, *p*=0.014), with lower concentrations associated with greater numbers of the ε4 allele. Phenotypic differences in Aβ_1–42_ levels were non-significant (F(1,173)=1.1, *p*=0.29). Co-pathology status (*p*=0.22), age (*p*=0.99), sex (*p*=0.77), and MMSE (*p*=0.86) were likewise not significantly associated with Aβ_1–42_ levels. Omnibus results when restricting the cohort to patients with negligible co-pathology also showed the effect of pathology type was significant (F(1,77)=32.0, *p*=2.49 × 10^−7^) while the effect of phenotype was not (*p*=0.88); all other effects were non-significant including the association with APOE (all *p*>0.31).

**Figure 1.**
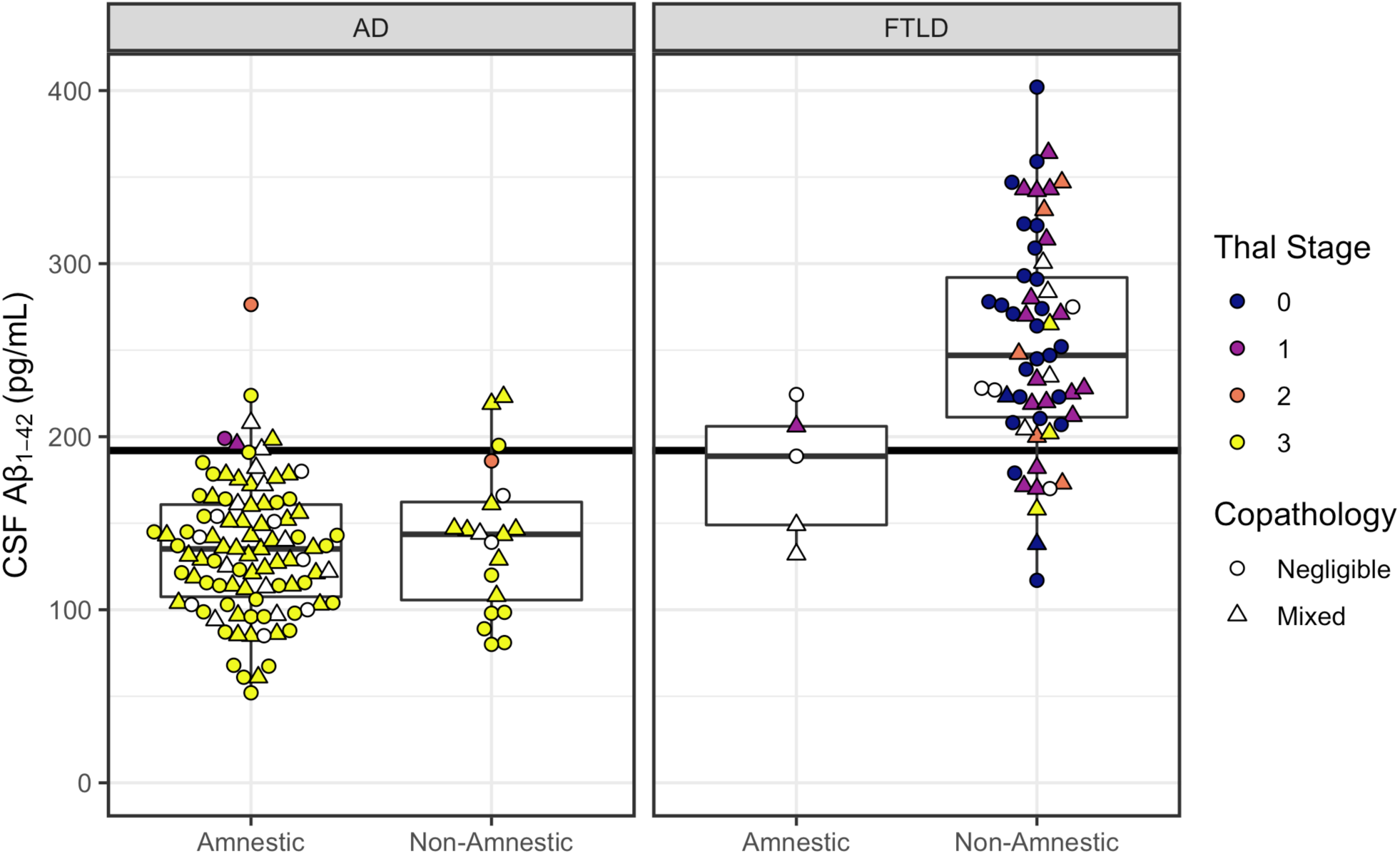
CSF *Aβ*_*1–42*_ concentrations according to phenotype (amnestic, non-amnestic) and postmortem ratings of amyloid-*β* pathology. Left panel plots patients with primary AD pathology. Right panel plots patients with primary FTLD pathology. Horizontal line indicates CSF Aβ_1–42_ = 192 pg/mL, with patients below the line designated as A+ status. Color indicates modified Thal staging of amyloid-β pathology on a zero-to-three scale (white indicates cases with no available Thal staging). Shape indicates presence or absence of co-pathologies.

### A Status

False-negative rates for A status were low for both amnestic (seven of 98) and non-amnestic (three of 20) patients with primary AD pathology, with Fisher’s test showing no difference across phenotype (*p*=0.37). In the patients with primary FTLD pathology, 12 of 64 were CSF A-positive (see Supplementary Material).

### Between-group comparisons of CSF p-tau

CSF p-tau levels were compared parametrically across pathology and phenotype (Figure 2). A Type II ANCOVA showed that p-tau levels differed by pathology (F(1,173)=28.3, *p*=3.29 × 10^−7^) and memory phenotype (F(1,173)=7.2, *p*=0.0081), with APOE status (*p*=0.45), co-pathology status (*p*=0.71), age (F(1,173)=4.2, *p*=0.043), sex (*p*=0.18), and MMSE (*p*=0.09) included as covariates. Post-hoc linear contrasts confirmed that the pathologic FTLD group had lower CSF p-tau than the pathologic AD group (t(173)=-5.3, *p*=3.23 × 10^−7^), and that non-amnestic AD patients had lower CSF p-tau than amnestic AD patients (t(173)=-3.2, *p=*0.0072). Non-amnestic FTLD patients had lower CSF p-tau than both amnestic AD (t(173)=-9.1, *p=*1.67 × 10^−15^) and non-amnestic AD (t(173)=-3.5, *p=*0.0031) patients. Amnestic FTLD patients had lower CSF p-tau than amnestic AD (t(173)=-4.5, *p=*1.23 × 10^−4^); no other comparisons were significant (*p*>0.11). Similar results were obtained when restricting the cohort to patients with negligible secondary pathology: we observed significant effects of pathology type (F(1,77)=24.2, *p*=4.77 × 10^−6^), phenotype (F(1,77)=6.0, *p*=0.016), and age (F(1,78)=10.3, *p*=0.0020). There was a marginal effect of MMSE (F(1,77)=3.4, p=0.070), and all other effects were non-significant (all *p*>0.65), and.

**Figure 2.**
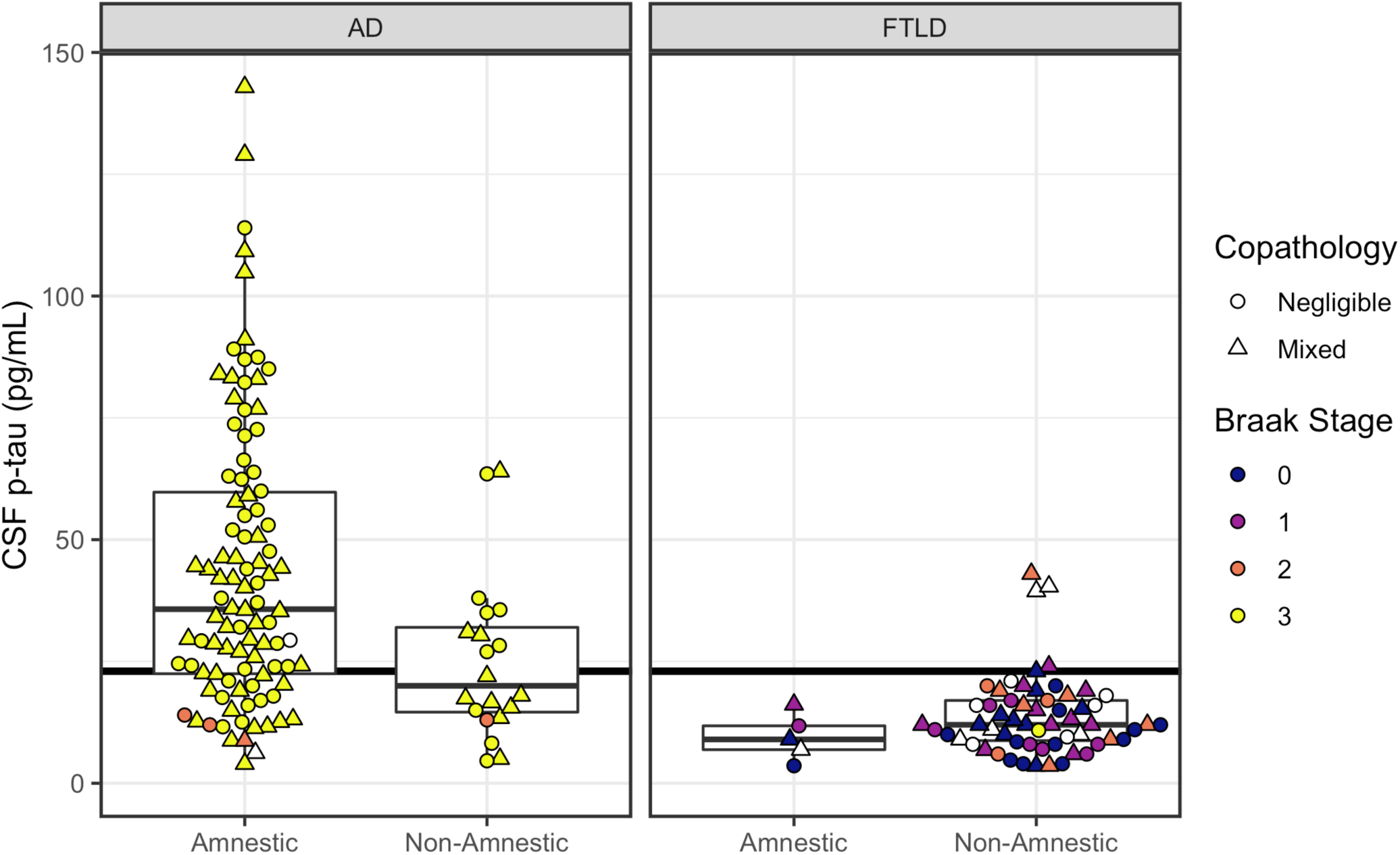
CSF p-tau concentrations according to phenotype (amnestic, non-amnestic) and postmortem ratings of tau neurofibrillary tangle pathology. Left panel plots patients with primary AD pathology. Right panel plots patients with primary FTLD pathology. Horizontal line indicates CSF p-tau = 23 pg/mL, with patients at or above the line designated as T+ status. Color indicates postmortem Braak staging of tau tangles (white indicates cases with no available Braak staging). Shape indicates presence or absence of co-pathology.

### T Status

Fisher’s test showed significantly lower rates for T-positivity in non-amnestic patients (eleven of 20; 50%) compared to amnestic patients (26 of 98; 24%) with confirmed AD pathology (OR = 0.30; CI: 0.10–0.89; *p*=0.018). Five of 64 FTLD patients were T-positive (Supplementary Material).

### Between-group comparisons of CSF t-tau

CSF t-tau levels were compared parametrically across pathology and phenotype (Figure 3). A Type II ANCOVA showed that CSF t-tau differed by pathology (F(1,173)=8.8, *p*=0.0034), phenotype (F(1,173)=9.1, *p*=0.0029), with APOE status (*p*=0.95), co-pathology status (*p*=0.14), age (*p*=0.44), sex (F(1,173)=11.0, *p*=0.0011), and MMSE (*p*=0.51) included as covariates. In post-hoc contrasts, the pathologic FTLD group had lower CSF t-tau than the AD group (t(173)=-3.0, *p*=0.0034). Compared to amnestic AD, we observed lower t-tau in non-amnestic AD (t(173)=-3.0, *p*=0.016) and in non-amnestic FTLD patients (t(173)=-6.6, *p*=2.98 × 10^−9^); all other comparisons were non-significant (*p*>0.19). CSF t-tau results were similar when restricting the cohort to patients with negligible co-pathology: the effect of pathology type was significant (F(1,77)=5.8, *p*=0.019), although the effect of memory phenotype was marginal (F(1,77)=3.5, *p*=0.063); all other effects were non-significant (all *p*>0.20).

**Figure 3.**
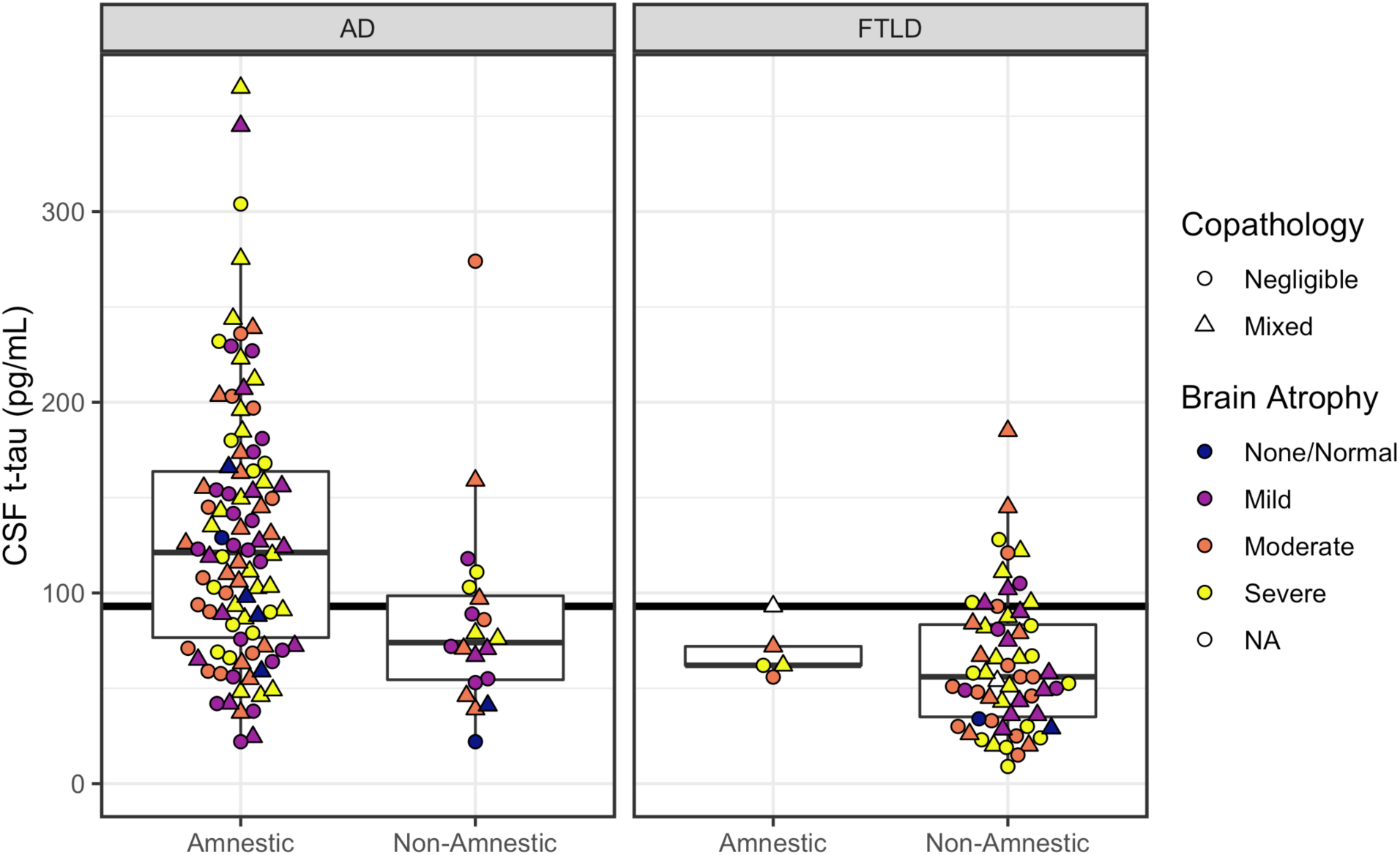
CSF t-tau concentrations according to phenotype (amnestic, non-amnestic) and postmortem ratings of brain atrophy. Left panel plots patients with primary AD pathology. Right panel plots patients with primary FTLD pathology. Horizontal line indicates CSF t-tau = 93 pg/mL, with patients at or above the line designated as N+ status. Color indicates severity of gross brain atrophy (white indicates cases with no available atrophy). Shape indicates presence/absence of co-pathology.

### N Status

Fisher’s test showed significantly lower rates of N-positivity for non-amnestic patients (six of 20, or 30%) compared to amnestic patients (65 of 98, or 66%) with confirmed AD pathology (OR = 0.22; CI: 0.06–0.68; *p*=0.005). Only 13 of 64 (20%) FTLD patients were positive for N. Two (3%) FTLD cases were true-negatives and had no atrophy at autopsy. The remaining 49 cases (77%) were likely false-negatives for N, showing mild to severe atrophy at autopsy (see Supplementary Material).

### Between-group comparison of CSF p-tau/Aβ_1–42_ levels

The p-tau/Aβ_1–42_ ratio was compared parametrically across pathology and phenotype (Figure 4). A Type II ANCOVA showed that p-tau/Aβ_1–42_ levels differed by pathology (F(1,173)=64.7, *p*=1.35 × 10^−13^) and memory phenotype (F(1,173)=8.5, *p*=0.0040), with APOE status (*p*=0.77), co-pathology status (*p*=0.95), age (F(1,173)=3.5, *p*=0.063), sex (*p*=0.27), and MMSE (*p*=0.14) included as covariates. Post-hoc testing confirmed that pathologic FTLD group had lower CSF t-tau than the AD group (t(173)=-8.0, *p*=1.35 × 10^−13^). Further, non-amnestic AD patients had significantly lower p-tau/Aβ_1–42_ ratios than amnestic AD patients (t(173)=-3.0, *p*=0.017). Non-amnestic FTLD patients had lower p-tau/Aβ_1–42_ ratios than both amnestic AD (t(173)=-12.2, *p*<0.00001) and non-amnestic AD (t(173)=-6.0, *p*=3.28 × 10^−8^); amnestic FTLD patients also had lower p-tau/Aβ_1–42_ ratios than both amnestic AD (t(173)=-4.9, *p*=1.26 × 10^−5^) and non-amnestic AD (t(173)=-2.7, *p*=0.031). No difference was observed between non-amnestic and amnestic FTLD patients (*p*=0.79). Omnibus results were highly similar when restricting the cohort to patients with no co-pathology: effects of pathology type (F(1,77)=50.7, p=4.91 × 10^−10^), phenotype (F(1,77)=4.5, *p*=0.037), age (F(1,78)=8.4, *p*=0.0050), and MMSE score (F(1,77)=4.2, *p*=0.044) were significant; and effects of sex and APOE ε4 allele count were non-significant (both *p*>0.76).

**Figure 4.**
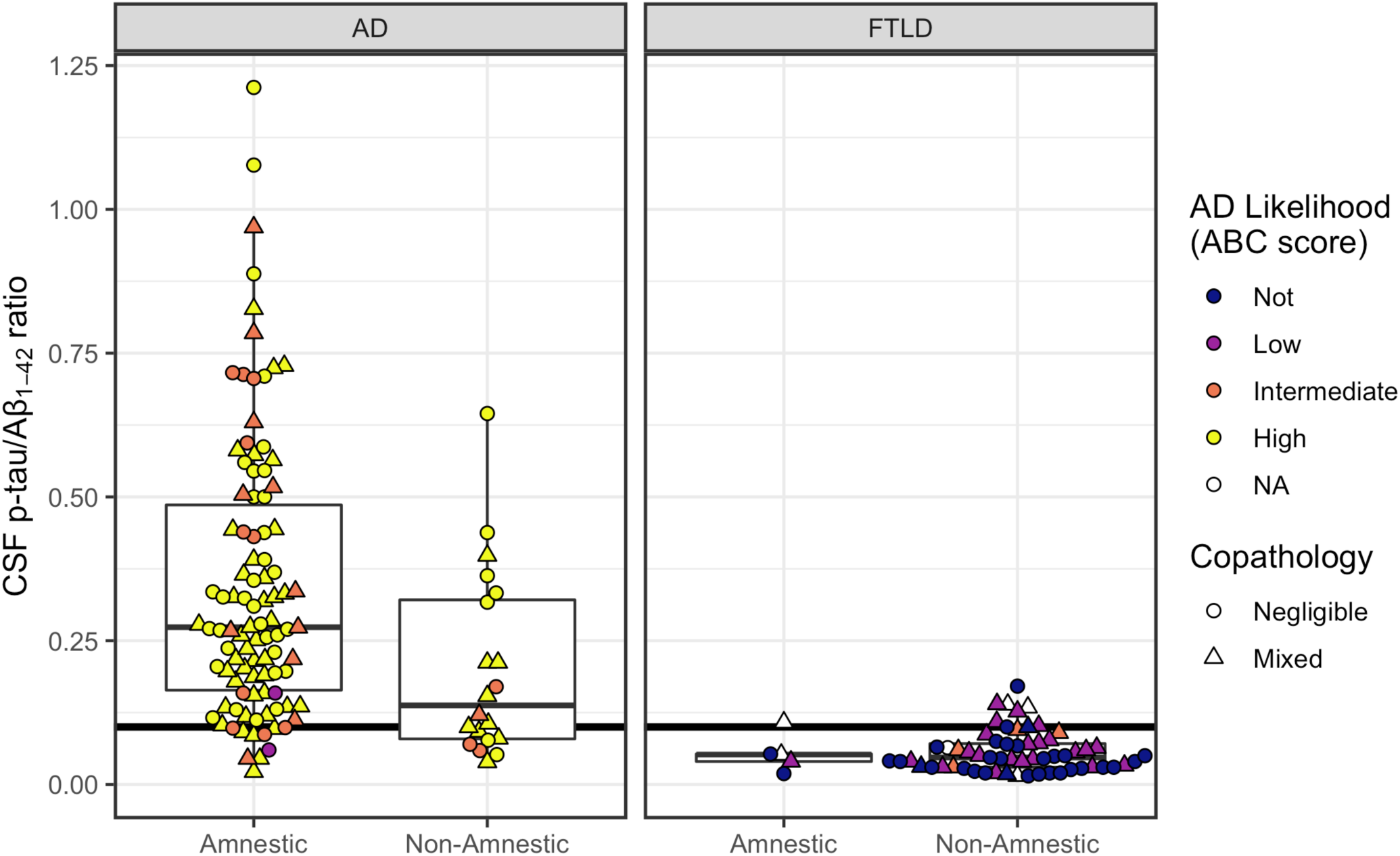
CSF p-tau/*Aβ*_*1–42*_ according to phenotype (amnestic, non-amnestic) and postmortem ratings of AD likelihood. Left panel plots patients with primary AD pathology. Right panel plots patients with primary FTLD pathology. Horizontal line indicates CSF p-tau/Aβ_1–42_ = 0.10, with patients at or above the line designated as positive for AD pathology. Color indicates level of AD pathologic change given by ABC score (white indicates cases with no available ABC score). Shape indicates presence/absence of co-pathology.

### AD status

A Fisher’s test showed significantly higher false-negative rates for non-amnestic patients (seven of 20, or 35%) compared to amnestic patients (10 of 98, or 10%) with confirmed AD pathology (OR = 4.7; CI: 1.27–16.58; *p*=0.009). In the pathologic FTLD group, 10 of 64 (16%) were AD positive (see Supplementary Material).

### Discrimination of pathologic AD from FTLD

We computed ROCs for each CSF measure stratifying amnestic or non-amnestic AD from all patients with FTLD pathology (Table 3). In an overall analysis distinguishing all AD from FTLD patients, CSF Aβ_1–42_ and the p-tau/Aβ_1–42_ ratio produced similar AUCs showing excellent discrimination of both markers (0.938 *vs*. 0.942). Of the two measures, CSF Aβ_1–42_ had the greatest sensitivity, while the p-tau/Aβ_1–42_ ratio yielded higher specificity (i.e., fewer primary FTLD patients classified as AD) than all other markers.

**Table 3:** ROC results for stratification of patients with AD from FTLD pathology. Includes area under the curve (AUC), best threshold or cut-point, and sensitivity and specificity at best threshold for each CSF measure. A) Stratification of all patients with AD from all patients with FTLD pathology. B) Stratification of amnestic patients with AD pathology from FTLD pathology. C) Stratification of non-amnestic patients with AD pathology from all FTLD pathology.

| **A) All AD from FTLD** |  |  |  |  |
| --- | --- | --- | --- | --- |
| CSF Measure | AUC | Threshold | Sensitivity | Specificity |
| $A\beta_{1-42}$ | 0.938 | 199.5 (pg/mL) | 0.958 | 0.812 |
| p-tau | 0.863 | 20.1 (pg/mL) | 0.737 | 0.906 |
| t-tau | 0.792 | 96.0 (pg/mL) | 0.585 | 0.875 |
| p-tau/$A\beta_{1-42}$ | 0.942 | 0.110 | 0.831 | 0.922 |
| **B) Amnesic AD from FTLD** |  |  |  |  |
| CSF Measure | AUC | Threshold | Sensitivity | Specificity |
| $A\beta_{1-42}$ | 0.941 | 199.5 (pg/mL) | 0.969 | 0.812 |
| p-tau | 0.888 | 20.1 (pg/mL) | 0.786 | 0.906 |
| t-tau | 0.822 | 96.5 (pg/mL) | 0.643 | 0.875 |
| p-tau/$A\beta_{1-42}$ | 0.958 | 0.110 | 0.888 | 0.922 |
| **C) Non-amnesic with AD from FTLD** |  |  |  |  |
| CSF Measure | AUC | Threshold | Sensitivity | Specificity |
| $A\beta_{1-42}$ | 0.925 | 168.0 (pg/mL) | 0.80 | 0.922 |
| p-tau | 0.743 | 12.5 (pg/mL) | 0.85 | 0.578 |
| t-tau | 0.647 | 66.5 (pg/mL) | 0.70 | 0.625 |
| p-tau/$A\beta_{1-42}$ | 0.866 | 0.076 | 0.80 | 0.781 |

DeLong’s test for two ROC curves compared diagnostic accuracy by AD phenotype for each marker. No difference in accuracy was observed between non-amnestic and amnestic AD when Aβ_1–42_ was used to discriminate groups from pathologic FTLD (*p*=0.64), which maintained high accuracy for both amnestic and non-amnestic AD (AUC=0.94 *vs*. AUC=0.92). However, CSF p-tau (D(105.1)=-1.9, *p*=0.057) and the p-tau/Aβ_1–42_ ratio (D(99.6)=-1.9, *p*=0.059) were both marginally less accurate for discriminating non-amnestic AD than amnestic AD from all FTLD patients. Although CSF t-tau is not a diagnostic marker, DeLong’s test shows a significantly lower AUC for non-amnestic AD than amnestic AD patients when discriminating from all FTLD patients (D(119.9)=-2.3, *p*=0.025).

For discrimination of amnestic AD from all FTLD patients, optimal cutoffs in the current study were close to the thresholds published by Shaw et al. (2009): for Aβ_1–42_, 199.5 pg/mL *vs*. 192 pg/mL; for p-tau, 20.1 pg/mL *vs*. 23 pg/mL; for t-tau, 96.5 pg/mL *vs*. 93 pg/mL; and for the p-tau/Aβ_1–42_ ratio, 0.11 *vs*. 0.10. However, for discrimination of non-amnestic AD from all FTLD patients, optimal cutoffs were substantially lower than published thresholds for Aβ_1–42_ (168.0 pg/mL), p-tau (12.5 pg/mL), t-tau (66.5 pg/mL), and the p-tau/Aβ_1–42_ ratio (0.076). ROC analyses were repeated excluding cases with co-pathology and produced similar results, although the p-tau/Aβ_1–42_ ratio had a high AUC (0.93) when discriminating non-amnestic AD from FTLD patients, albeit with a much lower cutoff of 0.051 (*vs*. 0.10) (Supplementary Table 2).

### Secondary analysis of p-tau associations with age and sex

In the current study, non-amnestic AD patients were younger at age of symptom onset and at death than amnestic AD patients, consistent with previous reports that atypical AD phenotypes are associated with a younger age of onset (Lam *et al.*, 2013; Mendez, 2017). While there was no effect for age in analyzing Aβ_1–42_ or t-tau, this demographic difference raised the question of whether lower p-tau values in the non-amnestic AD sample were simply a consequence of their younger age, which would be reflected by a positive association between age and p-tau levels among non-amnestic AD patients. In addition, models revealed significantly lower t-tau in males than females. Secondary analyses revealed that differences in CSF p-tau and t-tau between amnestic and non-amnestic AD patients could not be explained by age or sex differences between the groups (see Supplementary Material).

## Discussion

Recent diagnostic strategies, such as the ATN framework, have emphasized biomarkers over traditional clinical evaluations to improve antemortem predictions of AD pathology, as well as stratification from other pathologies such as FTLD. Even with the improved accuracy of antemortem predictions of AD pathology, subtle differences in CSF levels across AD variants might lead to a small number of diagnostic errors (Teng *et al.*, 2014; Paterson *et al.*, 2015; Wellington *et al.*, 2018; Pillai *et al.*, 2019). To better characterize the full spectrum of CSF profiles in AD continuum disease and evaluate diagnostic accuracy of CSF biomarkers, we compared the accuracy of CSF-based ATN classifications in AD patients with amnestic and non-amnestic clinical phenotypes, and in FTLD. In autopsy-confirmed primary AD pathology, non-amnestic AD patients were less likely than amnestic AD patients to be classified as A+T+N+, but more likely to be classified as normal (A-T-N-) according to CSF biomarkers. Parametric comparisons revealed that differences in designation are likely because non-amnestic AD patients have lower concentrations of both p-tau and t-tau in their CSF than amnestic AD. Likewise, ROC curve analyses showed that CSF p-tau and the p-tau/Aβ_1–42_ ratio have somewhat reduced AUC and lower sensitivity in non-amnestic AD than amnestic AD when discriminating from FTLD patients. While t-tau is not a diagnostic marker, it also had a lower AUC when discriminating non-amnestic AD from FTLD, emphasizing that concentrations of t-tau differ by amnestic phenotype. Unlike CSF p-tau and t-tau, CSF Aβ_1–42_ levels were equivalent between amnestic and non-amnestic AD patients, and were significantly lower in all FTLD patients. ROC curve analyses confirmed that CSF Aβ_1–42_ is a highly sensitive marker to the presence of AD in both amnestic and non-amnestic AD, and is excellent at stratifying both AD groups from FTLD patients. Thus, our findings indicate that A status is consistent across memory phenotypes in AD, and accurately detects individuals who are likely positive for pathologic amyloid deposition and thus AD continuum disease. However, more specific designations of T and N status based on CSF p-tau and t-tau may inaccurately classify non-amnestic AD patients as negative for tau aggregation or neurodegeneration. These findings highlight the need for careful implementation of the ATN framework for *in vivo* diagnosis, as markers optimized for the most common amnestic presentations of AD may not capture the full phenotypic spectrum of the pathologic disease.

While a primary goal of the ATN framework is to identify patients with AD pathology, a secondary goal of the ATN framework may be the characterization of individuals with non-AD pathology (Jack *et al.*, 2018). Under the ATN framework, detection of non-AD pathology relies on N-positive status in the absence of either A or T, or T-positive status in the absence of A. Our results show that patients with FTLD pathology were not reliably classified by ATN; only 12 of 64 were identified as having non-AD pathology (A-T+N-, A-T-N+, A-T+N+). Three FTLD patients were identified as having suspected co-occurring AD and non-AD pathologies; however two of these had ABC scores of 0 at autopsy. Instead, a majority of FTLD patients were N-negative due to lower CSF t-tau levels, despite showing atrophy at autopsy (only two cases of FTLD were judged not to be atrophic); nine of 64 were classified as having Alzheimer’s pathologic change, and the majority were classified as having normal biomarkers (40 of 64). It is noteworthy that a majority of non-amnestic AD patients were also N-negative (14 of 20). Discrepancies between t-tau results and autopsy assessments of atrophy in non-amnestic AD and in FTLD may indicate that brain structure was relatively preserved at time of lumbar puncture; alternatively, negative t-tau results may indicate a failure to detect true degeneration. While CSF t-tau is interpreted as a non-specific marker of neurodegeneration (Jack *et al.*, 2016), our results indicate that t-tau may be relatively insensitive to neurodegeneration in non-amnestic pathologic cases, both AD and FTLD.

We additionally tested discrimination based on p-tau/Aβ_1–42_ ratio as an alternative to the ATN framework. This composite marker had the highest AUC (0.96) and highest specificity (0.92) when stratifying amnestic AD from pathologic FTLD, a finding in agreement with previous studies (Vergallo *et al.*, 2017; Lleó *et al.*, 2018). Indeed, this ratio score correctly identified 90% of amnestic AD patients (Table 2). Results using the ATN framework were comparable: 93% of amnestic AD patients were classified as being on the AD continuum due to A-positive status. However, the p-tau/Aβ_1–42_ ratio had a lower AUC than Aβ_1–42_ alone (0.87 *vs*. 0.93) when discriminating non-amnestic AD patients from all FTLD patients, correctly identifying only 65% cases. By comparison, ATN designated 85% of non-amnestic AD patients as being on the AD continuum, although this increase was not significant.

The optimal thresholds for CSF Aβ_1–42_, p-tau, and p-tau/Aβ_1–42_ specific to our entire cohort were broadly consistent with cut-points that were established in an independent cohort to distinguish AD from healthy controls (Shaw *et al.*, 2009). However, all optimal cutoffs were lower when stratifying non-amnestic AD patients from FTLD, including CSF Aβ_1–42_. A caveat to this statement is that CSF t-tau is not intended to discriminate pathologic AD from pathologic FTLD; to calculate a true cutoff for t-tau in non-amnestic patients, we would need a reference control sample without degeneration, which is beyond the scope of this study. Still, lower optimal cutoffs of p-tau, t-tau, and the p-tau/Aβ_1–42_ reflect the lower concentrations of p-tau and t-tau in non-amnestic AD compared to amnestic AD. Even so, the sensitivity and specificity of these adjusted cutoffs is poor, and does little to improve diagnostic specificity of these markers due to overlap with pathologic FTLD. The exception is CSF Aβ_1–42_ which maintains high sensitivity and specificity in non-amnestic AD. Unlike p-tau and t-tau, the lower optimal cutoff of CSF Aβ_1–_ 42 in non-amnestic AD is a more conservative cut-off than in the entire sample, and may in part be an artifact of small cohort size; changing the cutoff from 192 to 166 would change the designation of only 1 subject (see Figure 1).

Instead of adjusting CSF cutoffs, our findings suggest that selection of the appropriate marker based on phenotype (amnestic or non-amnestic) and diagnostic goals (increase sensitivity or specificity) would do more to improve diagnostic accuracy. The p-tau/Aβ_1–42_ ratio appears to perform best in a cohort of amnestic AD dementia patients, and has slightly higher specificity for excluding non-AD cases. However, it performs less well in non-amnestic AD due to their lower levels of CSF p-tau. By comparison, the ATN framework has excellent sensitivity to the presence of Alzheimer’s continuum disease due in part to the reliable performance of A status as measured by Aβ_1–42_ in both amnestic and non-amnestic AD. However, it may be that more specific designations within the ATN framework incorrectly classify non-amnestic AD patients as normal or Alzheimer’s pathologic change. Potential limitations of classification using the ATN framework in non-amnestic AD should be considered for candidate treatment trials. Furthermore, our results indicate that the ATN framework is relatively insensitive to non-AD diseases like FTLD, and these patients were likely to be erroneously classified as having normal biomarkers or AD pathologic change alone. Biomarkers suggestive of co-occurring pathologies (A+T-N+) were unreliable: A+T-N+ failed to capture the majority of FTLD with secondary AD pathology and two of three A+T-N+ FTLD cases had no detected level of AD (zero ABC scores). In sum, both the ATN framework and the p-tau/Aβ_1–42_ were excellent at identifying amnestic AD cases, while the single marker of CSF Aβ_1–42_ had the most consistent performance across memory phenotypes in AD, and was sensitive to both amnestic and non-amnestic AD.

The current study additionally adds to a growing body of results on sex differences in CSF. Significantly lower levels of t-tau were observed in men compared to women with FTLD, and marginally lower levels of t-tau in men compared to women with AD (see Supplementary Material). Likewise, previous research has found that females have elevated CSF t-tau (Hohman *et al.*, 2018), a stronger association of *APOE-*ε4 with p-tau and t-tau (Hohman *et al.*, 2018), and higher AD pathologic burden and tau tangle density (Oveisgharan *et al.*, 2018) compared to males. Even so, phenotypic differences in CSF biomarkers could not be explained by group differences in global cognition or sex. Like sex, analysis of age effects did not support the interpretation that non-amnestic AD patients had less tau accumulation due to their younger age. On the contrary, younger patients (both amnestic and non-amnestic) in the pathologic AD group were likely to have higher levels of p-tau than older patients, in agreement with previous findings (Lleó *et al.*, 2019).

### Caveats and Limitations

There are a number of caveats to the current study. First, although autopsy studies suggest that mixed pathology is highly common in dementia, no current CSF analytes are able to determine the presence of co-occurring or “mixed” pathology. To increase the relevance and application of our findings, we examined patients who had AD and FTLD with negligible levels of secondary pathology thought to not contribute significantly to phenotype, as well as those with mixed pathology, including AD and FTLD patients with co-occurring α-synuclein pathology. To ensure that mixed pathology cases were not driving diagnostic errors, analyses included co-pathology status as a covariate and were repeated in patients with negligible co-pathology. Results were consistent across models with and without co-pathology cases, indicating that differences observed across phenotypes in AD are not explained by co-pathological status. We were insufficiently powered to examine subgroups of secondary pathology or subgroups of FTLD pathology. Second, while we interpret A-negative or T-negative status in AD as likely false-negative cases, and while the accumulation of these histopathologic features is thought to begin much earlier in the presymptomatic course of disease, we cannot entirely rule out the unlikely possibility that CSF markers accurately reflect a true absence of amyloid or tau pathology that developed only later in disease course and after obtaining the CSF sample. Third, amnestic and non-amnestic patients differed in age-related factors (age at CSF, at onset, and survival). These differences are expected given the dementia population: amnestic patients tend to be older at onset than non-amnestic variants of both AD and FTLD (Koedam *et al.*, 2010; Mendez, 2017). Still, one possible explanation of the higher incidence of T-negative and/or N-negative status in non-amnestic AD patients could be an earlier point in the course of disease at the time of CSF collection than amnestic AD. However, this was not supported by our data. To account for potential differences in the course of disease at the time of lumbar puncture, age at CSF and MMSE were included as covariates in models, and secondary analyses show CSF differences are not due to younger age in the non-amnestic AD group than AD. Even so, MMSE and other non-specific measures of disease severity may not capture differences across groups (Santana *et al.*, 2016; Pillai *et al.*, 2019). Fourth, t-tau has been criticized as a marker of neurodegeneration, in part because t-tau is highly correlated with p-tau, and thus characterizations of T-status and N-status may not be truly independent (Hampel *et al.*, 2018). Moreover, recent longitudinal work shows little prognostic value to t-tau in controls (Soldan *et al.*, 2019; Vos and Duara, 2019). While imaging-based markers of neurodegeneration may provide an attractive alternative to CSF t-tau, patients in the current study were not selected based on availability of MRI or PET data; and variability in the temporal intervals between patients’ scans and lumbar punctures, coupled with differences in scan acquisition parameters, might have reduced statistical power and injected noise into analysis results. However, we note that the shortcomings of t-tau as a marker of N are orthogonal to the phenotypic differences in ATN classification reported in the current study. Finally, this is a single-center study, and thus may be limited in generalizability.

In conclusion, results show that only CSF Aβ_1–42_ is consistent across amnestic and non-amnestic patients with AD pathology, and is highly accurate at discriminating AD pathology from FTLD in both memory groups. Even so, CSF Aβ_1–42_ has decreased specificity compared to other biomarkers. By comparison, CSF p-tau and t-tau levels differ across amnestic and non-amnestic patients with AD pathology, and show a much lower accuracy for non-amnestic patients. Because of these differences, more specific ATN designations within the Alzheimer’s continuum may be less accurate for non-amnestic variants of AD. Findings also show that the ATN framework had poor sensitivity to detecting primary FTLD, erroneously classifying them as Alzheimer’s pathologic change alone or as having normal biomarkers. Together, our results indicate that sensitivity of AD diagnoses can be improved by considering non-amnestic phenotype when making CSF-based designations of ATN status.

## Funding

This work was supported in part by NIH grants AG061277, AG054519, AG017586, and NS088341; Alzheimer’s Association grant AARF-16-443681; BrightFocus Foundation A2016244S; and the Penn Institute on Aging.

> KAQ Cousins is a Fellow for Advancing Research and Treatment for Frontotemporal Lobar Degeneration (ARTFL) and a recipient of the Alzheimer’s Association Research Fellowship. Jeffrey S. Phillips is the recipient of an Alzheimer’s Association Research fellowship and an NIA-sponsored K01 Mentored Research Scientist Career Development Award.

## Competing Interests

Authors have no conflicts of interest to report.

## Supplementary Material

### A status in FTLD

In the patients with primary FTLD pathology, 12 of 64 were CSF A-positive (three amnestic; nine non-amnestic). Of the 12 A-positive FTLD patients, three had definitive histopathologic assessments of no AD pathology (*i.e*., ABC scores all equal to zero), and two had CERAD scores of zero (missing Thal); these five cases thus represent false-positives and likely false-positives. Five A-positive cases had non-zero Thal staging (one, n=3; two, n=1, three, n=1); A-positive CSF in this context may accurately capture the co-occurrence of amyloid pathology.

### T status in FTLD

Of the five T-positive FTLD cases (five non-amnestic; one amnestic), two had postmortem Braak stage of zero and one, respectively; CSF p-tau results in these cases may thus represent false-positives. Braak staging was not performed for the remaining three T-positive cases in the FTLD group.

### N Status in FTLD

Of 13 N-positive cases within the pathologic FTLD group (12 non-amnestic; one amnestic), one lacked an atrophy rating, three had mild atrophy, four had moderate atrophy, and five had severe atrophy. Of the 51 (80%) N-negative FTLD cases, only two were judged at autopsy to have no atrophy; 11 had mild atrophy, 18 had moderate atrophy, 19 had severe atrophy, and one lacked an atrophy rating.

### AD status in FTLD determined by p-tau/Aβ_1–42_

In the pathologic FTLD group, 10 patients had supra-threshold p-tau/Aβ_1–42_ ratios. Three of 10 were definite false-positives and had zero ABC scores, two without co-pathology detected at autopsy and one with co-occurring hippocampal sclerosis. Four had low ABC scores, and another one with missing ABC scores had AD co-pathology according to pathologist’s judgment. The final two were missing ABC scores, but had co-occurring primary age-related tauopathy according to pathologist’s judgment.

### Secondary analysis of p-tau associations with age, and t-tau associations with sex

We performed a follow-up regression analysis with p-tau values for patients in the pathologic AD group, including a term for the interaction of age and phenotype. In the amnestic AD group, age was negatively associated with p-tau levels (β=-0.021, t=-2.9, *p*=0.0051); the corresponding effect of age on CSF p-tau in the non-amnestic group did not differ from the reference value, evidenced by a non-significant interaction (*p*=0.28) (Supplementary Figure 1). These results indicate that the younger average age of non-amnestic AD patients did not account for the lower p-tau values observed in that group. Instead, lower age was associated with higher CSF p-tau levels in AD.

Additionally, we performed secondary analyses to clarify sex-related effects on t-tau, as revealed by models. Patient groups were well-balanced in terms of sex distributions (Table 1; Supplementary Figure 2). We performed a follow-up regression analysis with t-tau values for patients in the pathologic AD group, including a term for the interaction of sex and phenotype. In the amnestic AD group, male sex was marginally associated with lower t-tau levels (β=-0.22, t= -1.95, *p*=0.053); the corresponding effect of sex on CSF t-tau in the non-amnestic group did not differ from the reference value, evidenced by a non-significant interaction (*p*=0.86) (Supplementary Figure 2). These results indicate that lower levels of t-tau in non-amnestic AD compared to AD cannot be attributed to differences in sex distribution.

## SUPPLEMENTARY FIGURES

**Supplementary Figure 1.**
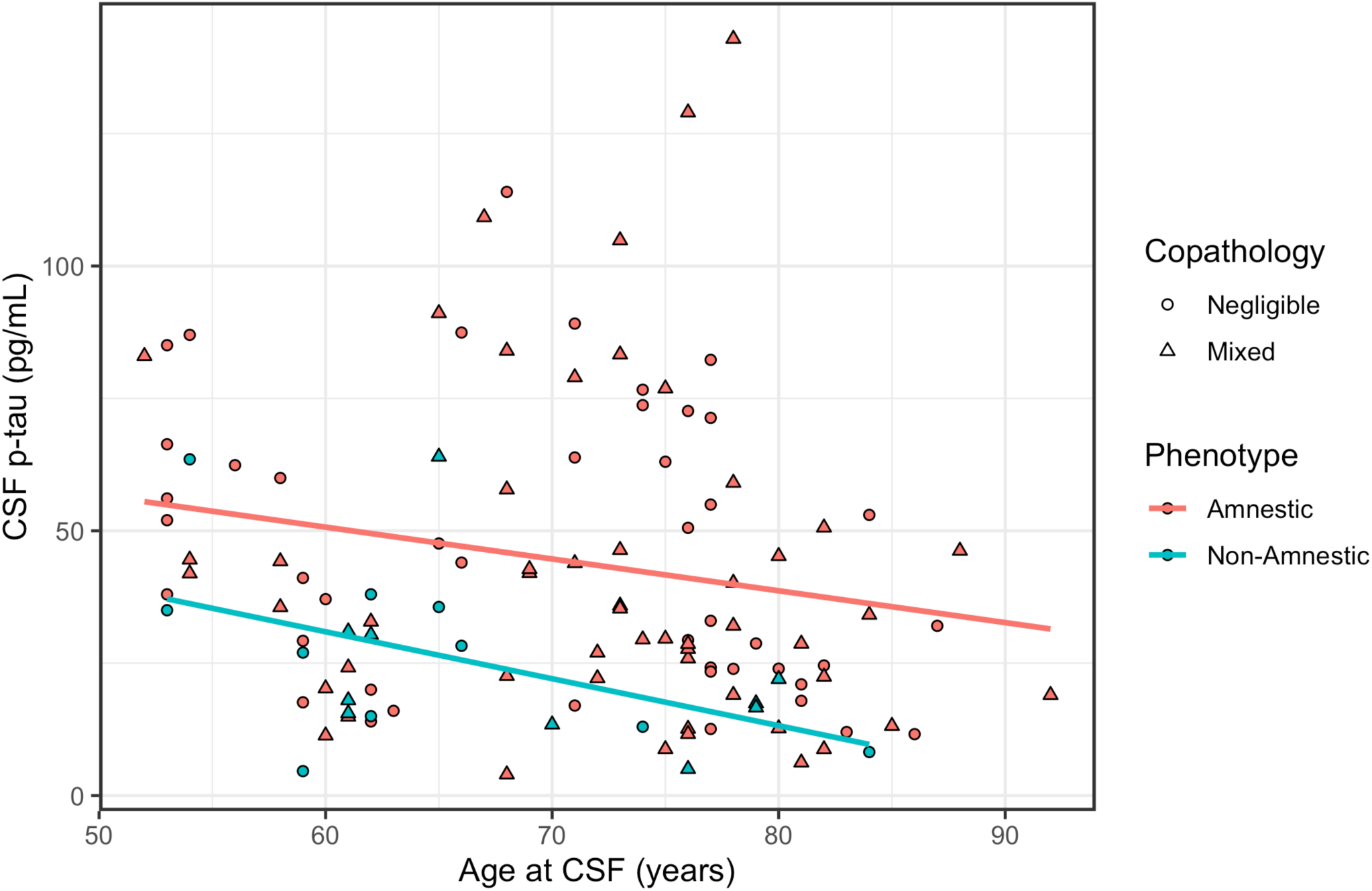
Cross-sectional associations of cerebrospinal fluid biomarkers with age among patients with autopsy-confirmed Alzheimer’s disease. Top panel plots values for CSF Aβ_1–42_, middle panel plots values for CSF p-tau, and bottom panel plots values for CSF t-tau by age. Color indicates phenotype. Shape indicates presence/absence of co-pathology.

**Supplementary Figure 2.**
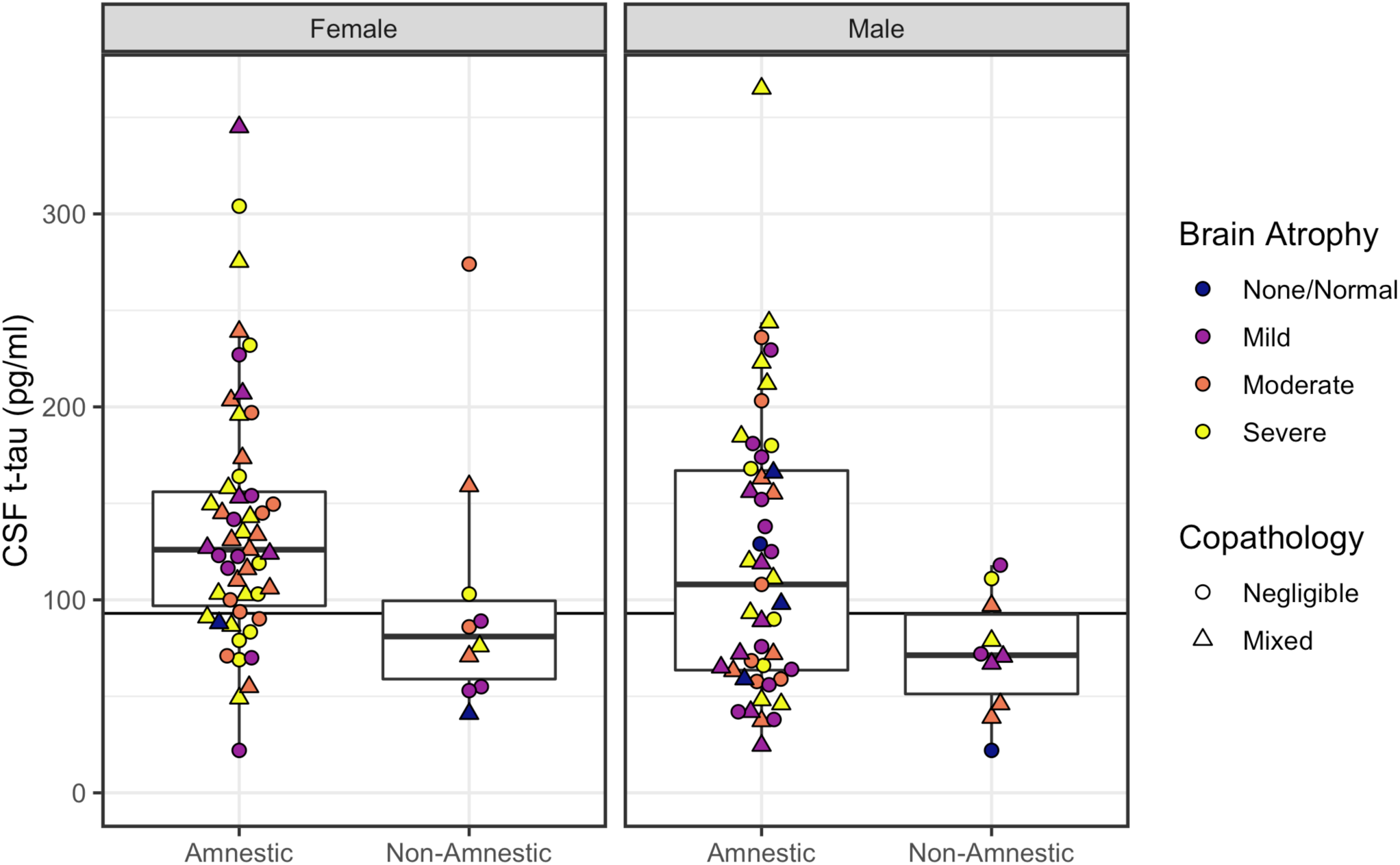
CSF p-tau concentration plotted according to phenotype and sex. Left panel plots patients with primary AD pathology; right panel plots patients with primary FTLD pathology. Horizontal line indicates CSF p-tau = 23 pg/mL, with patients at or above the line designated as T+ status. Color indicates severity of gross brain atrophy. Shape indicates presence/absence of co-pathology.

**Supplementary Table 1:** Results of ATN classification using CSF biomarkers, excluding cases with secondary pathology. Interpretation of biomarkers using ATN framework (above the black line) and p-tau/Aβ_1–42_ ratio (below the black line). Alzheimer’s disease, AD and co-occurring non-AD pathologic change, and AD pathologic change designations are considered Alzheimer’s continuum disease.

| Interpretation |  | Biomarker profile | Amnesic AD (n=45) | Non-amnesic AD (n=10) | Amnesic FTLN (n=2) | Non-amnesic FTLN (n=28) |
| --- | --- | --- | --- | --- | --- | --- |
| Alzheimer's continuum | Alzheimer's disease | A+T+N+ | 24 (53%) | 3 (30%) | 0 (0%) | 0 (0%) |
|  |  | A+T+N- | 9 (20%) | 3 (30%) | 0 (0%) | 0 (0%) |
|  | Alzheimer's pathologic change | A+T-N- | 6 (13%) | 2 (20%) | 1 (50%) | 1 (4%) |
|  | AD and concomitant suspected non-AD pathologic change | A+T-N+ | 3 (7%) | 1 (10%) | 0 (0%) | 2 (7%) |
| Non-Alzheimer's pathologic change |  | A-T+N- | 1 (2%) | 0 (0%) | 0 (0%) | 0 (0%) |
|  |  | A-T-N+ | 0 (0%) | 0 (0%) | 0 (0%) | 3 (11%) |
|  |  | A-T+N+ | 1 (2%) | 0 (0%) | 0 (0%) | 0 (0%) |
| Normal biomarkers |  | A-T-N- | 1 (2%) | 1 (10%) | 1 (50%) | 22 (79%) |
| Alzheimer's disease | | p-tau/A$\beta_{1-42}$ $\geq 0.1$ | 41 (91%) | 6 (60%) | 0 (0%) | 2 (7%) |

**Supplementary Table 2:** ROC results for stratification of patients with AD from FTLD pathology, excluding cases with secondary pathology. Includes area under the curve (AUC), best threshold or cut-point, and sensitivity and specificity at best threshold for each CSF measure. A) Stratification of all patients with AD from all patients with FTLD pathology. B) Stratification of amnestic patients with AD pathology from FTLD pathology. C) Stratification of non-amnestic patients with AD pathology from all FTLD pathology.

|  |  |  |  |  |
| --- | --- | --- | --- | --- |
| **A) All AD from FTLD** |  |  |  |  |
| CSF Measure | AUC | Threshold | Sensitivity | Specificity |
| $A\beta_{1-42}$<br>p-tau<br>t-tau | | | | |
| p-tau/$A\beta_{1-42}$ | | | | |
| **B) Amnesic AD from FTLD** |  |  |  |  |
| CSF Measure | AUC | Threshold | Sensitivity | Specificity |
| $A\beta_{1-42}$<br>p-tau<br>t-tau | | | | |
| p-tau/$A\beta_{1-42}$ | | | | |
| **C) Non-amnesic with AD from FTLD** |  |  |  |  |
| CSF Measure | AUC | Threshold | Sensitivity | Specificity |
| $A\beta_{1-42}$<br>p-tau<br>t-tau | | | | |
| p-tau/$A\beta_{1-42}$ | | | | |

## References

Andreasen N, Sjögren M, Blennow K. CSF markers for Alzheimer’s disease: total tau, phosphotau and Aβ42. world J Biol psychiatry 2003; 4: 147–155.

Armstrong MJ, Litvan I, Lang AE, Bak TH, Bhatia KP, Borroni B, et al. Criteria for the diagnosis of corticobasal degeneration. Neurology 2013; 80: 496–503.

Crutch SJ, Lehmann M, Schott JM, Rabinovici GD, Rossor MN, Fox NC. Posterior cortical atrophy. Lancet Neurol 2012; 11: 170–178.

Dickerson BC, McGinnis SM, Xia C, Price BH, Atri A, Murray ME, et al. Approach to atypical Alzheimer’s disease and case studies of the major subtypes. CNS Spectr 2017; 22: 439–449.

Ewers M, Mattsson N, Minthon L, Molinuevo JL, Antonell A, Popp J, et al. CSF biomarkers for the differential diagnosis of Alzheimer’s disease: a large-scale international multicenter study. Alzheimer’s Dement 2015; 11: 1306–1315.

Folstein MF, Folstein SE, McHugh PR. ‘Mini-mental state’. A practical method for grading the cognitive state of patients for the clinician. J Psychiatr Res 1975; 12: 189–198.

Galton CJ, Patterson K, Xuereb JH, Hodges JR. Atypical and typical presentations of Alzheimer’s disease: a clinical, neuropsychological, neuroimaging and pathological study of 13 cases. Brain 2000; 123: 484–498.

Gibbons GS, Banks RA, Kim B, Changolkar L, Riddle DM, Leight SN, et al. Detection of Alzheimer disease (AD)-specific tau pathology in AD and nonAD tauopathies by immunohistochemistry with novel conformation-selective tau antibodies. J Neuropathol Exp Neurol 2018; 77: 216–228.

Gibbons GS, Kim SJ, Robinson JL, Changolkar L, Irwin DJ, Shaw LM, et al. Detection of Alzheimer’s disease (AD) specific tau pathology with conformation-selective anti-tau monoclonal antibody in co-morbid frontotemporal lobar degeneration-tau (FTLD-tau). Acta Neuropathol Commun 2019; 7: 34.

Gorno-Tempini ML, Hillis AE, Weintraub S, Kertesz A, Mendez MF, Cappa SF, et al. Classification of primary progressive aphasia and its variants. Neurology 2011; 76: 1006–1014.

Hampel H, Toschi N, Baldacci F, Zetterberg H, Blennow K, Kilimann I, et al. Alzheimer’s disease biomarker-guided diagnostic workflow using the added value of six combined cerebrospinal fluid candidates: Aβ1–42, total-tau, phosphorylated-tau, NFL, neurogranin, and YKL-40. Alzheimer’s Dement 2018; 14: 492–501.

Hodges JR, Davies RR, Xuereb JH, Casey B, Broe M, Bak TH, et al. Clinicopathological correlates in frontotemporal dementia. Ann Neurol 2004; 56: 399–406.

Höglinger GU, Respondek G, Stamelou M, Kurz C, Josephs KA, Lang AE, et al. Clinical diagnosis of progressive supranuclear palsy: the movement disorder society criteria. Mov Disord 2017; 32: 853–864.

Hohman TJ, Dumitrescu L, Barnes LL, Thambisetty M, Beecham G, Kunkle B, et al. Sexspecific association of apolipoprotein e with cerebrospinal fluid levels of tau. JAMA Neurol 2018; 75: 989–998.

Igaz LM, Kwong LK, Xu Y, Truax AC, Uryu K, Neumann M, et al. Enrichment of C-terminal fragments in TAR DNA-binding protein-43 cytoplasmic inclusions in brain but not in spinal cord of frontotemporal lobar degeneration and amyotrophic lateral sclerosis. Am J Pathol 2008; 173: 182–194.

Jack CR, Bennett DA, Blennow K, Carrillo MC, Dunn B, Haeberlein SB, et al. NIA-AA Research Framework: Toward a biological definition of Alzheimer’s disease. Alzheimer’s Dement 2018; 14: 535–562.

Jack CR, Bennett DA, Blennow K, Carrillo MC, Feldman HH, Frisoni GB, et al. A/T/N: an unbiased descriptive classification scheme for Alzheimer disease biomarkers. Neurology 2016; 87: 539–547.

Jack CR, Knopman DS, Jagust WJ, Petersen RC, Weiner MW, Aisen PS, et al. Tracking pathophysiological processes in Alzheimer’s disease: An updated hypothetical model of dynamic biomarkers. Lancet Neurol 2013; 12: 207–216.

Koedam ELGE, Lauffer V, van der Vlies AE, van der Flier WM, Scheltens P, Pijnenburg YAL. Early-versus late-onset Alzheimer’s disease: more than age alone. J Alzheimer’s Dis 2010; 19: 1401–1408.

Lam B, Masellis M, Freedman M, Stuss DT, Black SE. Clinical, imaging, and pathological heterogeneity of the Alzheimer’s disease syndrome. Alzheimers Res Ther 2013; 5: 1.

Lleó A, Alcolea D, Martínez-Lage P, Scheltens P, Parnetti L, Poirier J, et al. Longitudinal cerebrospinal fluid biomarker trajectories along the Alzheimer’s disease continuum in the BIOMARKAPD study. Alzheimer’s Dement 2019; 15: 742–753.

Lleó A, Irwin DJ, Illán-Gala I, Mcmillan CT, Wolk DA, Lee EB, et al. A 2-Step Cerebrospinal Algorithm for the Selection of Frontotemporal Lobar Degeneration Subtypes. JAMA Neurol 2018; 75: 738–745.

Mackenzie IRA, Neumann M, Bigio EH, Cairns NJ, Alafuzoff I, Kril J, et al. Nomenclature and nosology for neuropathologic subtypes of frontotemporal lobar degeneration: an update. Acta Neuropathol 2010; 119: 1.

McKeith IG, Dickson DW, Lowe J, Emre M, O’Brien JT, Feldman H, et al. Diagnosis and management of dementia with Lewy bodies: Third report of the DLB consortium. Neurology 2005; 65: 1863–1872.

McKhann GM, Knopman DS, Chertkow H, Hyman BT, Jack CR, Kawas CH, et al. The diagnosis of dementia due to Alzheimer’s disease: Recommendations from the National Institute on Aging-Alzheimer’s Association workgroups on diagnostic guidelines for Alzheimer’s disease. Alzheimer’s Dement J Alzheimer’s Assoc 2011; 7: 263–269.

Mendez MF. Early-onset Alzheimer disease. Neurol Clin 2017; 35: 263–281.

Mirra SS, Heyman A, McKeel D, Sumi SM, Crain BJ, Brownlee LM, et al. The Consortium to Establish a Registry for Alzheimer’s Disease (CERAD): Part II. Standardization of the neuropathologic assessment of Alzheimer’s disease. Neurology 1991; 41: 479.

Montine TJ, Phelps CH, Beach TG, Bigio EH, Cairns NJ, Dickson DW, et al. National institute on aging-Alzheimer’s association guidelines for the neuropathologic assessment of Alzheimer’s disease: A practical approach. Acta Neuropathol 2012; 123: 1–11.

Murray ME, Graff-Radford NR, Ross OA, Petersen RC, Duara R, Dickson DW. Neuropathologically defined subtypes of Alzheimer’s disease with distinct clinical characteristics: a retrospective study. Lancet Neurol 2011; 10: 785–796.

Oeckl P, Steinacker P, Feneberg E, Otto M. Neurochemical biomarkers in the diagnosis of frontotemporal lobar degeneration: an update. J Neurochem 2016; 138: 184–192.

Onyike CU, Diehl-Schmid J. The epidemiology of frontotemporal dementia. Int Rev Psychiatry 2013; 25: 130–137.

Oveisgharan S, Arvanitakis Z, Yu L, Farfel J, Schneider JA, Bennett DA. Sex differences in Alzheimer’s disease and common neuropathologies of aging. Acta Neuropathol 2018; 136: 887–900.

Palmqvist S, Zetterberg H, Mattsson N, Johansson P, Minthon L, Blennow K, et al. Detailed comparison of amyloid PET and CSF biomarkers for identifying early Alzheimer disease. Neurology 2015; 85: 1240–1249.

Paterson RW, Toombs J, Slattery CF, Nicholas JM, Andreasson U, Magdalinou NK, et al. Dissecting IWG-2 typical and atypical Alzheimer’s disease: insights from cerebrospinal fluid analysis. J Neurol 2015; 262: 2722–2730.

Perry DC, Miller BL. Frontotemporal dementia. In: Seminars in neurology. Thieme Medical Publishers; 2013. p. 336–341

Phillips JS, Das SR, McMillan CT, Irwin DJ, Roll EE, Da Re F, et al. Tau PET imaging predicts cognition in atypical variants of Alzheimer’s disease. Hum Brain Mapp 2018; 39: 691–708.

Phillips JS, Da Re F, Dratch L, Xie SX, Irwin DJ, McMillan CT, et al. Neocortical origin and progression of gray matter atrophy in nonamnestic Alzheimer’s disease. Neurobiol Aging 2018; 63: 75–87.

Phillips JS, Da Re F, Irwin DJ, McMillan CT, Vaishnavi SN, Xie SX, et al. Longitudinal progression of grey matter atrophy in non-amnestic Alzheimer’s disease. Brain 2019; 142: 1701–1722.

Pillai JA, Bonner-Jackson A, Bekris LM, Safar J, Bena J, Leverenz JB. Highly Elevated Cerebrospinal Fluid Total Tau Level Reflects Higher Likelihood of Non-Amnestic Subtype of Alzheimer’s Disease. J Alzheimer’s Dis 2019; 70: 1051–1058.

Pouclet-Courtemanche H, Nguyen T-B, Skrobala E, Boutoleau-Bretonnière C, Pasquier F, Bouaziz-Amar E, et al. Frontotemporal dementia is the leading cause of “true” A−/T+ profiles defined with Aβ42/40 ratio. Alzheimer’s Dement Diagnosis, Assess Dis Monit 2019; 11: 161–169.

Rascovsky K, Hodges JR, Knopman D, Mendez MF, Kramer JH, Neuhaus J, et al. Sensitivity of revised diagnostic criteria for the behavioural variant of frontotemporal dementia. Brain 2011; 134: 2456–2477.

Robinson JL, Lee EB, Xie SX, Rennert L, Suh E, Bredenberg C, et al. Neurodegenerative disease concomitant proteinopathies are prevalent, age-related and APOE4-associated. Brain 2018; 141: 2181–2193.

Santana I, Duro D, Lemos R, Costa V, Pereira M, Simões MR, et al. Mini-Mental State Examination: Screening and diagnosis of cognitive decline, using new normative data. Acta Med Port 2016; 29: 240–248.

Shaw LM, Vanderstichele H, Knapik-Czajka M, Clark CM, Aisen PS, Petersen RC, et al. Cerebrospinal fluid biomarker signature in alzheimer’s disease neuroimaging initiative subjects. Ann Neurol 2009; 65: 403–413.

Soldan A, Pettigrew C, Fagan AM, Schindler SE, Moghekar A, Fowler C, et al. ATN profiles among cognitively normal individuals and longitudinal cognitive outcomes. Neurology 2019; 92: e1567–e1579.

Strong MJ, Abrahams S, Goldstein LH, Woolley S, Mclaughlin P, Snowden J, et al. Amyotrophic lateral sclerosis-frontotemporal spectrum disorder (ALS-FTSD): Revised diagnostic criteria. Amyotroph lateral Scler Front Degener 2017; 18: 153–174.

Struyfs H, Niemantsverdriet E, Goossens J, Fransen E, Martin J-J, De Deyn PP, et al. Cerebrospinal fluid P-Tau181P: biomarker for improved differential dementia diagnosis. Front Neurol 2015; 6: 138.

Tapiola T, Alafuzoff I, Herukka S-K, Parkkinen L, Hartikainen P, Soininen H, et al. Cerebrospinal fluid β-amyloid 42 and tau proteins as biomarkers of Alzheimer-type pathologic changes in the brain. Arch Neurol 2009; 66: 382–389.

Teng E, Yamasaki TR, Tran M, Hsiao JJ, Sultzer DL, Mendez MF. Cerebrospinal fluid biomarkers in clinical subtypes of early-onset Alzheimer’s disease. Dement Geriatr Cogn Disord 2014; 37: 307–314.

Toledo JB, Brettschneider J, Grossman M, Arnold SE, Hu WT, Xie SX, et al. CSF biomarkers cutoffs: the importance of coincident neuropathological diseases. Acta Neuropathol 2012; 124: 23–35.

Toledo JB, Van Deerlin VM, Lee EB, Suh E, Baek Y, Robinson JL, et al. A platform for discovery: the University of Pennsylvania integrated neurodegenerative disease biobank. Alzheimer’s Dement 2014; 10: 477–484.

Vergallo A, Carlesi C, Pagni C, Giorgi FS, Baldacci F, Petrozzi L, et al. A single center study: Aβ42/p-Tau 181 CSF ratio to discriminate AD from FTD in clinical setting. Neurol Sci 2017; 38: 1791–1797.

Vos SJB, Duara R. The prognostic value of ATN Alzheimer biomarker profiles in cognitively normal individuals. 2019

Wellington H, Paterson RW, Suárez-González A, Poole T, Frost C, Sjöbom U, et al. CSF neurogranin or tau distinguish typical and atypical Alzheimer disease. Ann Clin Transl Neurol 2018; 5: 162–171.

Wolk DA. Amyloid imaging in atypical presentations of Alzheimer’s disease. Curr Neurol Neurosci Rep 2013; 13: 412.

Xie SX, Baek Y, Grossman M, Arnold SE, Karlawish J, Siderowf A, et al. Building an integrated neurodegenerative disease database at an academic health center. Alzheimer’s Dement 2011; 7: e84–e93.

